# Human intention inference with a large language model can enhance brain-computer interface control: A proof-of-concept study

**DOI:** 10.1101/2025.06.01.657282

**Authors:** Shue Shiinoki, Seitaro Iwama, Junichi Ushiba

**Author notes:** Correspondence Junichi Ushiba, Ph.D. Department of Biosciences and Informatics, Faculty of Science and Technology, Keio University, 3-14-1 Hiyoshi, Kouhoku-ku, Yokohama, Kanagawa 223-8522, Japan.

## Abstract

Brain–computer interface (BCI) control enables direct communication between the brain and external devices. However, the accuracy of BCI control based on intentions inferred from unstable neural signals remains limited, even with data-driven approaches to tailor neural decoders. Here we propose a knowledge-driven framework for inferring human intentions, leveraging large language models (LLMs) that incorporate prior knowledge about human behavior and neural activity. We developed a neural decoder that integrates neural and oculomotor signals with contextual information using an LLM agent. Its feasibility was tested in a real-world BCI task involving interaction with a computer application. The LLM-based decoder achieved an average accuracy of 79% among participants whose neural signals were responsive (11 out of 20) in inferring the intention to select arbitrary posts in a social networking service. Ablation analyses revealed that the integration of contextual information, multimodal signals, and empirical knowledge is critical for decoding accuracy. This study demonstrates the feasibility of a neural decoding framework using an LLM, paving the way for improved performance in BCI-driven external device operation for individuals with disabilities.

**Highlights:**

- Large language models can infer human intent by integrating neural and oculomotor signals with screen context.
- The proposed model outperforms conventional data-driven approaches in decoding accuracy.
- Ablation analyses reveal that integrating contextual information, multimodal signals, and empirical knowledge is critical for accurate decoding.

## Introduction

A brain-computer interface (BCI) capable of bypassing human motor pathways to express user volition has long been desired for augmented alternative communication^1–5^. To achieve this, a computational model, namely a neural decoder, is employed to infer user intentions embedded in real-time neural signals^6–8^. The decoder plays a critical role in a BCI system, as the decoded outputs, which are translated into device commands, need to be accurate enough to reconstruct user intentions. The accuracy from noninvasive modalities has been significantly improved over the decades owing to advances in statistical methods and machine learning algorithms^9,10^.

One representative approach to enhancing the BCI performance is deep learning, which enables data-driven representation learning from minimally preprocessed neural signals^8,11,12^. A variety of network structures, such as recurrent neural networks (RNNs) or convolutional neural networks (CNN), have been proposed to extract features from neural signals without relying on prior knowledge or assumptions about the biological model^13,14^. CNN-based models have consistently demonstrated high performance, leading to the development of derivative models aimed at improving accuracy, generalization across various tasks and subjects^8,11,14–19^.

Despite remarkable achievements of deep learning in fields such as image processing and speech recognition owning to its generalization ability and high accuracy^20–22^, such breakthroughs have not yet been realized in neural decoding, particularly when using low-fidelity neural measurements such as the scalp electroencephalogram (EEG) to read out of human cortical activity. This limitation likely stems from the short duration of experimental sessions, which makes it impractical to collect datasets that sufficiently cover the vast data space required to learn representations that generalize across datasets. These challenges highlight the need for methods that alleviate the fully data-driven nature of existing approaches.

An alternative to fully data-driven model training is to incorporate knowledge-driven signal observation. For instance, in a Bayesian framework, prior knowledge can be used for initiating training from a more informative probability distribution. A more flexible integration of such prior knowledge can be achieved by using a large language model (LLM) as an inference module. LLMs are suitable for scenarios with limited neural training data as they embed neurophysiological knowledge that can support decoding as a complement to the input signals. Indeed, LLMs have demonstrated human-level reasoning abilities in knowledge-driven applications such as recommendation systems^23^, lifelong learning^24^, and autonomous driving tasks^25,26^. In autonomous driving, for instance, LLM-based systems that mimic human-like decision-making based on common sense and past experience have achieved highly generalized performance through causal reasoning, surpassing conventional reinforcement learning methods^25,26^.

Here, we propose a multimodal decoding system that infers motor intentions and operational targets from EEG and eye movements, based on knowledge-driven reasoning using an LLM. We evaluated its effectiveness in BCI control within a real-world scenario. Participants used the BCI system while interacting with a social networking service to assess its feasibility. The performance of the LLM-based decoder was compared against variants with ablated inputs or knowledge, data-driven baselines, and analyses of classification outcomes. We hypothesized that the effectiveness of the LLM-based BCI decoder arises from the integration of multimodal signal observation, task-specific decoding instructions, and a small amount of participant-derived training data.

## Results

### Decoder architecture

In this proof-of-concept study, we propose a decoder framework that integrates EEG and oculomotor signals, contextual information, knowledge, and a small number of EEG training samples. These inputs are all transformed into text and fed into a large language model (LLM) agent during BCI operation (Fig. 1, Supplementary Movie). The LLM agent is instructed to reason which content displayed on the PC screen the user intended to select (Fig.1, Supplementary Movie). The LLM agent was instructed to reason out which content shown in the PC display the user selected.

**Fig. 1.**
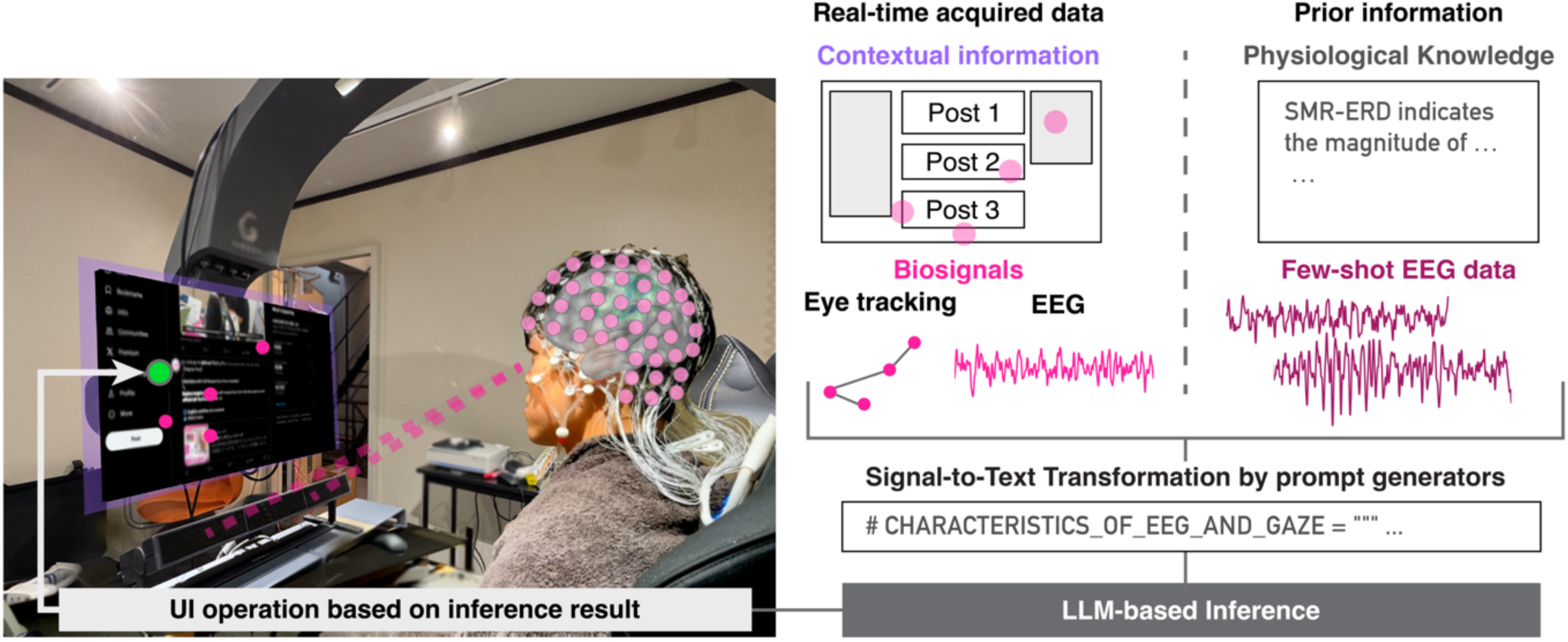
Schematic of real-time human intention inference using an LLM-based agent. Neural and oculomotor signals, namely, eye-tracking and scalp electroencephalogram (EEG) data—as well as contextual information (i.e., content displayed on the PC screen) were individually processed and transformed into text data at the initial stage of the processing pipeline. In parallel, prior information, including physiological knowledge from previous EEG decoding studies and a few samples of subject-specific EEG data, was also converted into text. This multimodal textual information was then input into the large language model (LLM) for inference. Based on the inference result, the PC cursor automatically moved to the location corresponding to the participant’s intended action.

All input data are converted into textual representations using specialized modules, allowing the LLM agent to interpret them in accordance with the given instructions. The agent employs a “chain-of-thought” reasoning process for inferring human intention, a methodology that has been successfully applied in other fields, such as autonomous driving. In this study, we adopt the ReAct framework^25,27^ to guide the reasoning process based on the LLM’s embedded common knowledge. This framework combines logical inference with information extracted from neural signals, along with physiological and heuristic knowledge provided as prompts. ReAct addresses the final decision-making problem (e.g., determining user attention or intent to press a button) through iterative steps of action selection, observation of outcomes, and reasoning based on those outcomes.

To support this process, the system leveraged multiple information sources as agent tools (i.e., candidate actions), including action intention references, EEG feature extraction results, the number of visual fixations within each tweet display area, and their spatial distribution. In addition, environmental data, such as tweet and button coordinates, UI metadata, and physiological knowledge, are incorporated as prior information (Fig. 1b). Prompts are generated by integrating text-based descriptions of EEG decoding knowledge (see Supplementary Information) and a few-shot EEG dataset collected before BCI operation. These prompts are then provided to the LLM agent, which performs reasoning in a structured format and outputs the corresponding operation commands.

### The proposed decoder is feasible to infer the user intention

Before the formal evaluation of decoder accuracy, the signal quality of EEG and eye gaze signals was visually inspected. For EEG, the temporal, spatial, and spectral information of EEG relevant to neural decoding was extracted using the spatio-spectral decomposition^28^ to visualize time-frequency representations (Supplementary Fig. 1). Specifically, the event-related desynchronization of sensorimotor rhythm (SMR-ERD) found during motor attempt was extracted. For the eye gaze signal, we visualized scan path maps indicating sequences of fixations on displays.

First, we investigated whether the proposed LLM-based decoder could perform neural decoding. Participants were stratified based on the responsiveness of their scalp EEG signals and grouped into responders and non-responders. Responsiveness was determined according to SMR-ERD magnitude (Fig. 2a). We evaluated decoding accuracy for both action and target selection (i.e., the “push like” or “read” command, and the tweet selection), as well as for action detection alone. The results showed that the responder group outperformed the non-responder group in both overall decoding and action decoding alone (Fig. 2b and Supplementary Fig. 2a; *t*-test, *p* < 0.05; Cohen’s *d* = 2.86, 2.94), suggesting that the proposed decoding approach is feasible and its performance is dependent on SMR-ERD responsiveness.

**Fig. 2.**
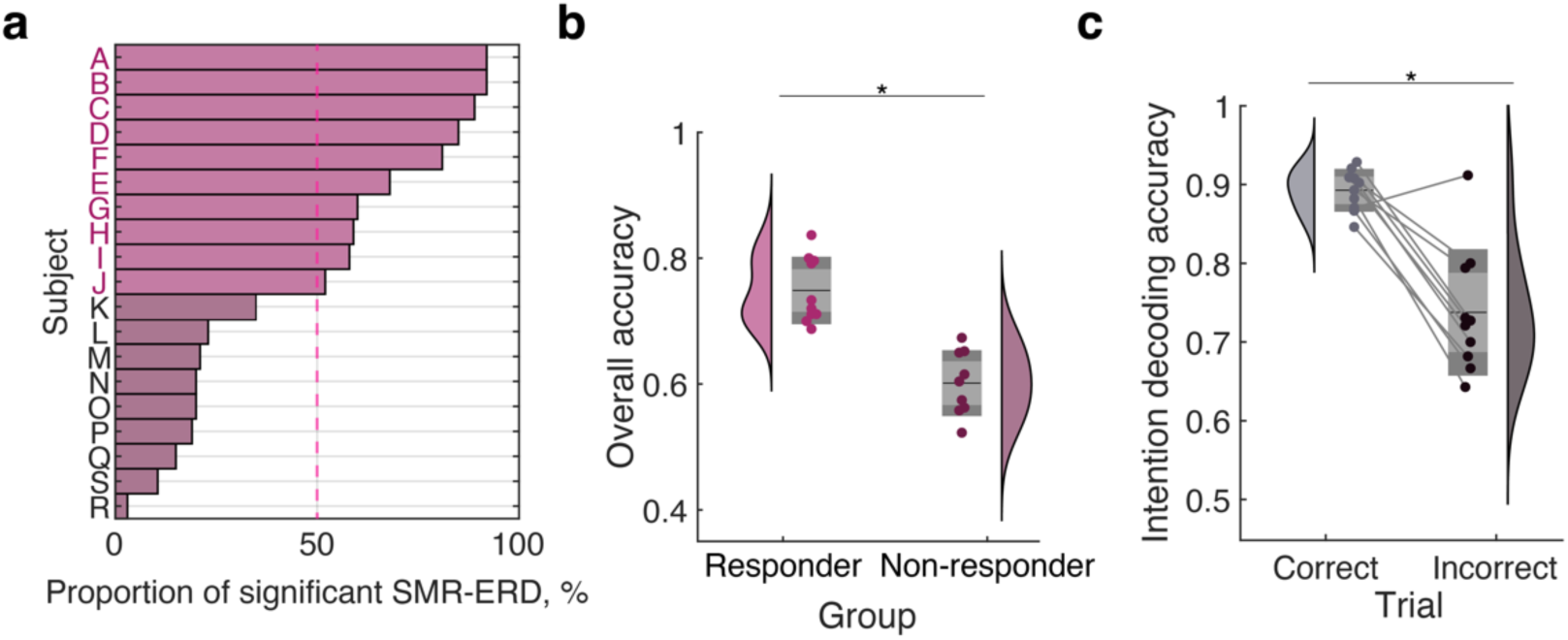
Evaluation of decoding accuracy during brain–computer interfacing. **a.** Participant-wise proportion of significant SMR-ERD in the first block. A threshold of 50% was used to categorize participants into responder and non-responder groups. Responders are highlighted. **b**. Comparison of overall decoding accuracy between responder and non-responder groups, based on intended action and target decoding. **c**. Comparison of intention decoding accuracy between trials in which the target was correctly versus incorrectly inferred.

Next, we examined how action decoding accuracy influences target decoding accuracy, in order to assess the interaction between the two decoding processes. To this end, target decoding accuracy was compared between trials in which motor intention was correctly versus incorrectly detected. We found that target decoding was significantly more accurate when motor intention was correctly detected (Fig. 2c; paired t-test, *p* < 0.05; Cohen’s *d* = 1.88). Furthermore, when target selection was accurate, action decoding accuracy was also significantly higher (Supplementary Fig. 2b; paired t-test, *p* < 0.05; Cohen’s *d* = 1.58). These findings suggest that successful motor intention decoding is associated with enhanced target decoding performance.

### Multimodal signal input, prior knowledge, and few-shot information are necessary for effective decoding

In the present study, multimodal data, namely neural and oculomotor movements, were used in an integrated manner for inferring human intention. We investigated the contribution of each modality to overall decoding performance. First, we conducted an ablation study on input modalities (Fig. 3a), modifying the prompt to perform the same inference task as the original model. A repeated measures analysis of variance (rmANOVA) revealed a significant decline in decoding accuracy when either signal was removed (main effect: *F* = 3.45, *p* < 0.05). Post-hoc pairwise *t*-tests showed that the decoder using both EEG and gaze signals (*EEG (+), Gaze (+)*) significantly outperformed the decoder using only EEG (*EEG (+), Gaze (−)*: *t* = 3.3, *p* < 0.05, Cohen’s *d* = 1.04) and the decoder using only gaze (*EEG (−), Gaze (+)*: *t* = 2.4, *p* < 0.05, Cohen’s *d* = 0.77). These results indicate that integrating both modalities significantly improves decoding performance over using either modality alone.

**Fig. 3.**
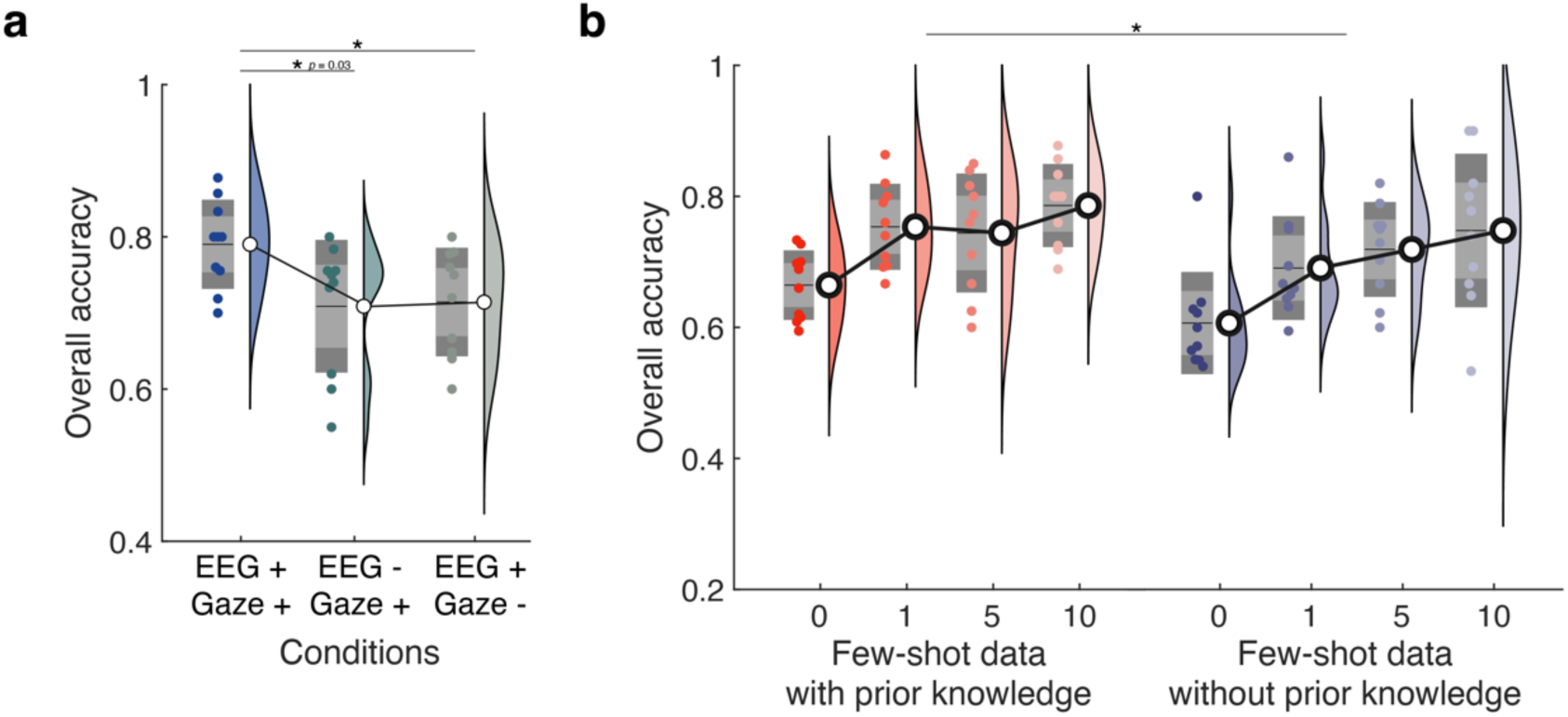
Multimodal signals and prior knowledge are necessary for inference s**a.** Accuracy of decoding based on multimodal and single-modal data conditions. **b.** Accuracy of decoding with prior knowledge and few-shot data.

Next, we examined the effects of two factors on decoding performance: empirical knowledge of EEG decoding, provided as prior information in the prompts, and few-shot EEG data from the initial trials, included as past decoding experience (Fig. 3b). To assess whether prior knowledge could compensate for limited data and whether decoding performance could be maintained with minimal examples, a two-way rmANOVA was conducted. The independent variables were the presence of empirical knowledge and the number of few-shot trials, while the dependent variable was decoding accuracy. The results showed significant main effects for both factors, but not interaction. Post-hoc tests indicated that including heuristic knowledge significantly improved performance in the 0-shot, 1-shot, 5-shot, and 10-shot conditions (Paired *t*-test, *p*<0.05, Cohen’s *d* = 1.08, 0.98, 0.55, 0.39, Bonferroni correction for multiple comparisons). Here, the number of “shots” refers to the number of trials provided, with each trial containing four sample data points. Furthermore, 10-shot decoding significantly outperformed both 0-shot and 5-shot conditions (*p* < 0.05, paired *t*-tests with Bonferroni correction; Cohen’s *d* = 1.84, 0.60, respectively).

### Comparison with other data-driven decoder architectures

The decoding performance of the knowledge-driven approach was evaluated in comparison with data-driven methods. Specifically, we compared the proposed method with CNNs and a Light Gradient Boosting Machine (LightGBM) ^29^. While the standard input for CNN-based EEG decoding is raw data, in this study, the CNN model was also provided with SMR-ERD features, identical to those used as input to the LLM.

We assessed decoding accuracy across multiple CNN models and the proposed model (Fig. 4a). The LLM decoder used 10 trials of participant-specific EEG data as few-shot prompts. We compared this with several configurations of EEGNet^17^ trained under varying data conditions: Fine-tuning (10 trials): Trained from scratch on 10 trials of participant-specific data, matching the quantity used in the LLM decoder. Fine-tuning (50 trials): Trained on 50 trials (i.e., 5 sessions) of data from the participant. Pretraining + Fine-tuning (pretrained + 10 trials): First pre-trained on 500 trials from other participants, then fine-tuned using 10 trials from the same participant as in the test. The 10-shot LLM decoder achieved significantly higher accuracy than both the CNN model trained on 5 sessions and the CNN model trained on 1 session (*p* < 0.05, paired *t*-test; Cohen’s *d* = 1.23, 1.09). These results suggest that the performance of the LLM decoder is comparable to that of a CNN model trained with pretraining and fine-tuning.

**Fig. 4.**
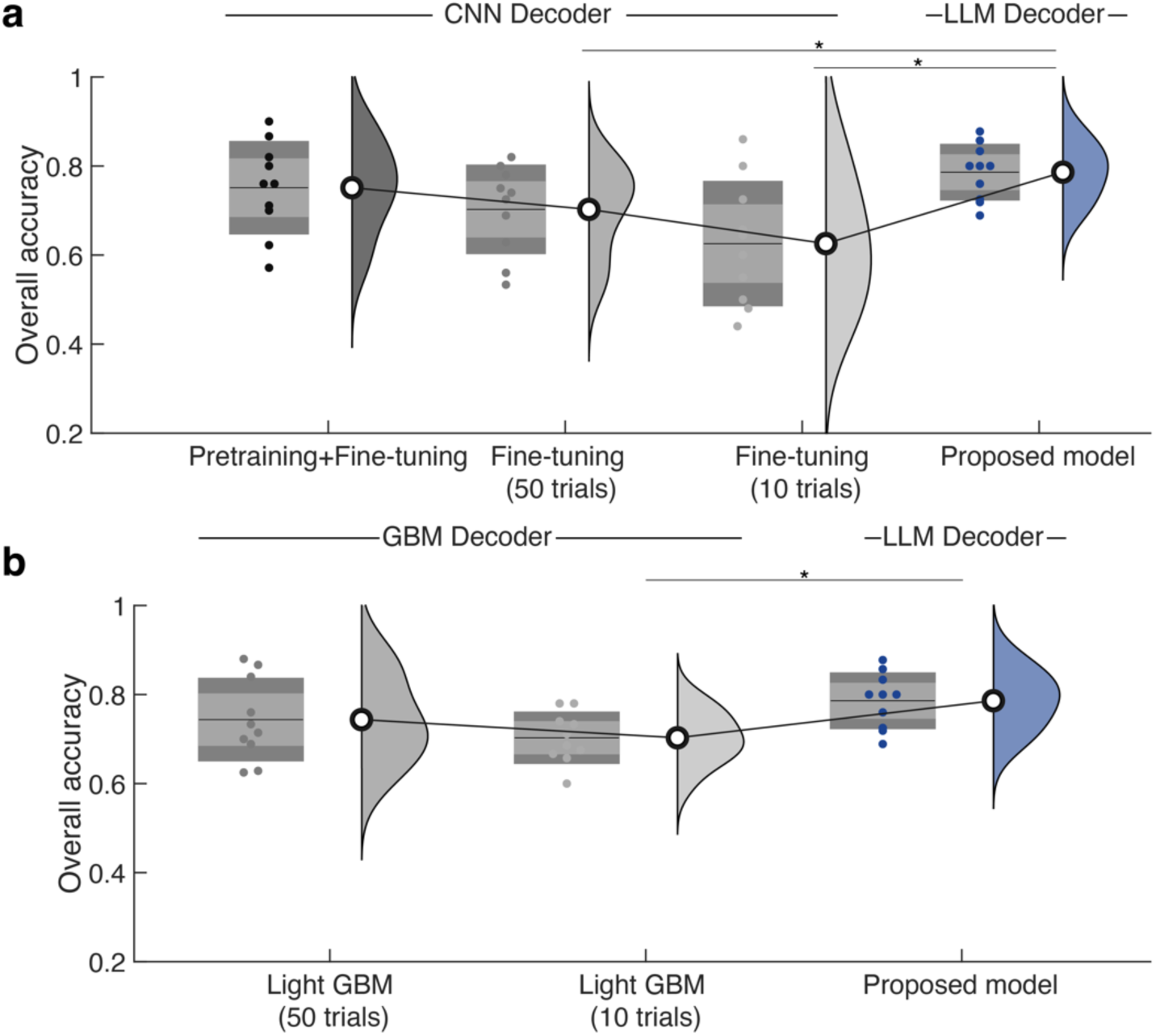
Comparisons of decoding accuracy with different decoder architectures. **a.** Decoding accuracy comparison with convolutional neural network (CNN) models. **b.** Decoding accuracy comparison with light gradient boosting machine (GBM) models.

To evaluate the effectiveness of the LLM approach compared to a decision tree-based method, we also trained a LightGBM model using the same SMR-ERD features as inputs. We then compared the decoding accuracy of the 10-shot LLM decoder to that of the LightGBM model trained on 5-session and 1-session datasets (Fig. 4b). The 10-shot LLM decoder was significantly more accurate than the 1-session LightGBM model (*p* < 0.05, paired *t*-test; Cohen’s *d* = 1.5). Collectively, these results indicate that our model can enhance BCI control accuracy under limited user data compared to the fully data-driven models.

## Discussion

In the present study, we investigated the feasibility of an LLM-based neural decoder architecture for inferring human movement intentions and their corresponding targets. Through performance evaluations and ablation analyses, we demonstrate that the proposed decoder effectively infers movement intentions and target selections by integrating neural and oculomotor signals with contextual and empirical knowledge.

Specifically, the decoder achieved significantly higher accuracy for responders, with successful motor intention decoding closely associated with improved target selection. Ablation analyses revealed that multimodal signal integration, empirical knowledge, and few-shot learning each contributed significantly to decoding performance, with knowledge-driven inference mitigating the limitations imposed by limited training data. These results support the feasibility of an LLM-driven approach for enhancing BCI control accuracy under a data-constrained conditions.

The limited amount of available data from individual users has critically hindered the development of high-performance EEG decoders. In contrast to fields such as computer vision and natural language processing where training datasets often exceed 10 million sample is typically in the more than 10 million scale^13,21^ and model architectures are correspondingly large in both parameter count and representational capacity, EEG-based deep learning models remain relatively small due to data scarcity, making it difficult to learn complex representations. Consequently, data-driven algorithms for EEG decoding have attempted to improve performance by reducing model complexity and employing regularization techniques^17,18^, but classification accuracy and generalization ability remain limited^11,14^. In contrast, the present LLM-based architecture enables researchers to incorporate decoding techniques into top-down inference through prompt design. In this study, we demonstrated that leveraging knowledge-driven inference via an LLM allows for accurate inference of motor intention using fewer than 50 trials of EEG data. With minimal training data and text-based descriptions of decoding strategies, our approach offers a promising alternative to conventional data-driven models, which have been constrained by limited dataset sizes in EEG decoding.

One benefit to use an LLM agent for neural decoding is to integrate information from multiple sources. Previous studies have attempted to augment the degrees of freedom in BCI control by independently utilizing multimodal signals such as eye gaze and EEG. This approach enables the simultaneous decoding of interaction targets and behavioral intentions based on gaze and EEG signals, respectively. However, treating these modalities separately may hinder the extraction of movement-related information from gaze signals, and vice versa^30–32^. Specifically, eye movements not only convey information about the object to be manipulated but also reflect behavioral intentions such as the user’s intent to interact with an object when fixating on it, or anticipation of upcoming tasks as indicated by gaze shifts^33–36^. The significantly higher decoding performance observed when combining EEG and eye movement signals, compared to using EEG alone, highlights the benefits of an integrated decoding approach. Moreover, data-driven methods face a challenge: integrating heterogeneous modalities within a single model typically requires increased model complexity, in accordance with the no-free-lunch theorem, thereby demanding larger training datasets. In contrast, LLMs can readily accommodate such multimodal integration when appropriate descriptors are constructed for each modality—leveraging prior knowledge and the generic, flexible input format of natural language, as demonstrated in the present study.

As demonstrated in the ablation analysis for LLM prompting (Fig. 3b), we found that decoding performance improves with few-shot prompting, even when physiological or empirical knowledge about EEG decoding is provided as input. If the performance degradation typically observed with a small number of few-shot examples is mitigated by the inclusion of empirical knowledge, this suggests that such knowledge compensates for decoding-relevant information that the LLM would otherwise extract from the data. This implies that empirical knowledge supplements but does not replaceinformation essential for decoding. However, our results indicate that few-shot prompting and empirical knowledge independently contribute to performance improvements. This suggests that, in addition to the physiological and empirical knowledge provided as text, the LLM also extracts useful decoding information from the few-shot data. One possible source of this information is the participant-specific characteristics of EEG signals, particularly in relation to SMR-ERD. Previous studies have shown that inter-participant variability in EEG features such as ERD magnitude and pattern variability constrains the performance of fully data-driven decoding approaches^37,38^. Our findings suggest that the LLM leverages these subject-specific ERD characteristics embedded in the few-shot input to enhance inferential performance.

Although the present study demonstrated the feasibility of performing knowledge-driven inference of behavioral intentions including motor intentions from a small amount of data using an LLM, several limitations remain. First, the generalization of performance across and within participants was limited by the ability to consistently elicit SMR-ERDs.

Currently, the inference system’s ability to accurately classify behavioral intentions is restricted to evaluations conducted during the best-performing sessions specifically, those in which participants with sufficient SMR-ERD reactivity were able to reliably generate SMR-ERDs throughout the session.

Second, although this inference system encodes all input information in text form for the LLM, recent advances in multimodal LLMs including models capable of processing image inputs have shown promise in domains such as GUI control^39–41^ and medical imaging^42^. In the present study, we restricted the input modality to text in order to isolate and evaluate the reasoning capability of the LLM without confounding effects from image-based input parsing. However, incorporating image input may be beneficial, as it aligns more closely with how human researchers interpret data through visual representation. Third, the decoding latency is relatively high, with an approximate delay of one minute. This delay primarily stems from the use of remote LLM inference via application programming interfaces, where both network communication and the LLM’s processing time contribute significantly to overall latency. We consider this a minor technical issue that could be resolved by deploying the LLM locally, thereby eliminating network overhead. Furthermore, ongoing advances in hardware and inference optimization (e.g., more efficient decoding algorithms) are expected to further reduce response times and mitigate the current latency bottleneck.

In summary, we demonstrated that a knowledge-based, LLM-driven neural decoding approach is feasible for BCI control under conditions of limited data availability. This framework enables experimenters to design task-specific prompts for inference, making it adaptable to various BCI paradigms beyond motor imagery-based BCIs. By leveraging prior physiological knowledge, the proposed approach has the potential to reduce calibration time without compromising decoding accuracy, representing a promising step toward practical applications in assistive technologies.

## Methods

### Participants

Twenty neurologically healthy adults (19 males, 1 female; mean age ± SD: 24 ± 1.4 years) were informed of the aim of the experiment and provided written informed consent. The experiment was approved by the local ethics committee of Keio University (IRB approval number: 2024-009) and was conducted in accordance with the Declaration of Helsinki. All participants were confirmed to be right-handed using the Edinburgh Handedness Inventory^43^.

### Experimental setup and protocol

EEG was recorded using a 128-channel HydroCel Geodesic Sensor Net from Electrical Geodesics Incorporated (EGI, Eugene, Oregon, USA) at a sampling rate of 1000 Hz. The impedance level of the EEG electrodes was maintained below 50 kΩ throughout the experiment. In this study, a PC-operated social media application, “X,” was used. “X” allows users to view short posts—commonly known as tweets—such as informational and news posts from celebrities and friends, and to rate these posts by “liking” them. A display showing the content list screen of “X” was placed in front of the participant. Eye movements were recorded at a sampling rate of 1200 Hz using a Tobii Pro Spectrum screen-based eye tracker (Tobii Technology AB, Danderyd, Stockholm, Sweden) positioned at the bottom of the display. The eye tracker was synchronized with the display to avoid measurement errors. Each participant performed the task of liking a tweet on the “X” GUI using motor imagery, while EEG and eye movements were simultaneously recorded using the behavioral intention decoding BCI described below.

The experiment consisted of one reference session and five evaluation sessions. After each session, participants completed the Quebec Assistive Technology Satisfaction Evaluation (QUEST) 2.0 questionnaire^44^ and a free-description questionnaire regarding their impressions of the BCI system to validate its usability. Participants were instructed to rest, read the on-screen content, and press the Like button using motor imagery during the time window shown in Fig. 1. During each trial, they were instructed to perform the task within the same screen without scrolling. To present a different UI layout in each trial, scrolling was performed at the end of the trial to refresh the content displayed. The decoder generated four inferences per trial, each based on five seconds of recorded data. Specifically, two inferences were made during the “read” phase and two during the “push like” phase. The inference results were presented via the GUI during the final “blank” period of each trial, and the participant manipulated the results using a cursor. The duration of the “blank” period was sufficient to accommodate inference processing, which took approximately 20–30 seconds.

### EEG Processing

EEG analysis was used to input EEG information into the decoding system. The recorded EEG signals were subjected to digital signal processing to remove artifacts. First, a third-order Butterworth band-stop filter (50 Hz) and a third-order Butterworth band-pass filter (3–70 Hz) were applied^45–47^. A spatial filtering technique called spatio-spectral decomposition (SSD) was then applied to the processed signals. SSD is capable of extracting neuronal oscillations with spatial patterns corresponding to SM1 activity^28^. The SSD algorithm extracted neural oscillations at the individual alpha frequency (IAF), which more effectively reflects the physiological properties of EEG than a uniform frequency band^48–50^. IAF was identified using the specparam algorithm^51^, which parameterizes and separates the periodic and aperiodic components of EEG signals and allows for the calculation of the frequency at which signal magnitude in the band of interest is maximal. The EEG input to the decoding system was the sensorimotor rhythm event-related desynchronization (SMR-ERD), which is thought to reflect sensorimotor excitability^52–54^.

SMR-ERD is defined as the relative change in power and is calculated as follows:

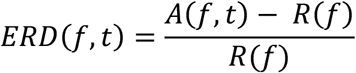

Where *A* represents the signal power of the EEG signals at time *t* and *f* represents the frequency, *R* and represents the mean signal power of the previous 9 s and recalculated in every 9 s (i.e., at the beginning of relaxation and imagery periods). The power was calculated using a short-time Fourier transform with a 90% overlap of the 1-second sliding window.

### Eye movement processing

To input eye movement information into the decoding system, we employed fixations to represent the positions on the screen where the user was gazing^55^. Fixations were extracted using an I-VT (Identification by Velocity Threshold) filter, which distinguishes between fixations and saccades based on eye movement velocity at each time point. A velocity threshold of 30°/s was applied^56^. Fixations were defined as sequences of gaze points in which velocity remained below this threshold for at least 100 ms. The resulting fixation points were mapped to predefined screen coordinates, and fixation clusters were further quantified for each tweet to determine gaze allocation across stimuli.

### Decoder Architecture

EEG signals, eye movements, environmental information in the form of the X application’s UI, and prior knowledge including EEG decoding expertise and past EEG data were integrated and input into an LLM to infer the user’s behavioral intentions. Within the signal descriptor, EEG, eye movement, and UI information were each preprocessed and converted into textual data. The specific processing procedures for each type of information are described below. Prompts were generated from the textual representations of the signal and environmental data, domain-specific knowledge regarding EEG decoding, and few-shot EEG data obtained during the reference session (see below). These prompts were then input into the LLM. The LLM performed inference based on a predefined format and ultimately output operation commands, which were used to control the application.

SMR-ERD time series data over a 5-second window were calculated from the EEG signals using the online EEG analysis described earlier. From this time series, candidate features were extracted. The feature extraction included the computation of various statistics, such as the mean, median, negative 1 SD, positive 1 SD, negative 2 SD, positive 2 SD, standard deviation, maximum amplitude, location of maximum amplitude, maximum and minimum values, peak-to-peak intervals, zero-crossing count, number of positive and negative values, and signal volatility. A gradient boosting model (LightGBM)^29^ was trained using preliminary EEG data to classify behavioral intention classes based on these features. Feature importance was then computed from the trained model, and the eight most informative features were selected for use. These included the mean, median, 1 SD, zero-crossing count, maximum amplitude value, location of maximum amplitude, percentage of negative values, and volatility.

Eye movement data were processed from the stable fixation points identified by the online analysis described earlier. UI information was also processed into a textual format that captured the user’s attentional focus. This was achieved by combining the two-dimensional spatial distribution of eye movements on the screen with the number of stable fixations per tweet. The resulting textual data included coordinates corresponding to the positions of tweets and the location of each tweet’s Like button.

To enable knowledge-driven reasoning, relevant knowledge for EEG decoding was also provided as input. This included physiological knowledge supporting the role of SMR- ED as a biomarker for SM1 excitability during motor imagery, heuristic insights into how EEG signals are affected by artifacts, and few-shot ERD data collected during the reference session.

### Decoder Evaluation

One of the 20 participants exhibited significant body motion artifacts in their EEG recordings, compromising signal quality. Therefore, this participant was excluded from the analysis. To objectively evaluate the performance of the decoder, the remaining 19 participants were grouped according to their responsiveness to SMR-ERD, and performance was evaluated separately for each group. First, a one-sample t-test was performed across trials for each participant’s ERD data at the individual alpha frequency (IAF) during the reference session. The null hypothesis assumed a mean ERD value of zero. Participants whose ERD values during the motor imagery period were significantly less than zero (i.e., negative direction) for more than 50% of the time interval were categorized as the responder group. Those who did not meet this criterion were categorized as the low responder group.

The performance of the decoder was evaluated based on the accuracy of decoding the presence or absence of the motor intentions “Read” and “Push Like,” as well as the accuracy of identifying the target tweets associated with these actions. Additionally, the overall decoding performance of the complete behavioral intention, including identification of the specific tweet that was the target of the action, was assessed. For the decoding of the complete behavioral intention, the self-reported responses indicating which tweets the participants “liked” at the end of each trial were used as the ground truth. In each trial, two datasets were obtained corresponding to “Push Like” and “Read.” Since no class imbalance was observed between these conditions, decoding performance was evaluated using overall accuracy.

For data-driven benchmarking, we implemented EEGNet^17^, a compact convolutional neural network (CNN) architecture optimized for EEG-based brain-computer interface (BCI) tasks. The model consisted of three convolutional layers with batch normalization and dropout, and was trained using the Adam optimizer with early stopping based on validation loss. We compared three training conditions: (1) 10-shot training using subject-specific data; (2) 5-session training using data from the same subject; and (3) pretraining on 500 trials from other subjects followed by fine-tuning on 10 trials from the target subject. All models used SMR-ERD features as input to ensure a fair comparison with the LLM-based decoder.

We compared the accuracy of the LLM-based decoder, which uses 10 trials of subject-specific data as few-shot examples within the prompt, with that of EEGNet models trained under different data conditions. Specifically, we prepared: (1) a model trained on 10 trials of data from the same subject—equivalent in size to the LLM’s few-shot input, (2) a model trained on 5 sessions (50 trials) of data from the same subject, and (3) a model pre-trained on 500 trials from other subjects and fine-tuned on 10 trials from the target subject. For all models, training was conducted for up to 200 epochs with early stopping based on validation loss. Training convergence was observed when the patience parameter for early stopping was set to 50.

To verify whether decoding by the LLM is superior to simple decision tree-based classification of individual features, we trained a LightGBM model to classify between the “Read” and “Push Like” motor intentions. The model was constructed and evaluated under the same training data conditions as those used for the CNN models. Early stopping was also employed, with training converging using a maximum of 2000 boosting rounds and an early stopping patience of 300 rounds.

### Statistical Analysis

All data were analyzed using parametric tests following verification of normality with the Shapiro–Wilk test. For multilevel datasets, a repeated-measures mixed ANOVA was conducted, followed by post hoc pairwise t-tests with Bonferroni correction. Sphericity was assessed using Mauchly’s test, and the Greenhouse–Geisser correction was applied when the assumption of sphericity was violated. Paired datasets were analyzed using paired t-tests.

## Conflicts of Interest

J.U. is a founder and representative director of the university startup company, LIFESCAPES Inc., involved in the research, development, and sales of rehabilitation devices, including brain-computer interfaces. SI and He receive a salary from LIFESCAPES Inc., and hold shares in LIFESCAPES Inc. This company does not have any relationships with the device or setup used in the current study. The remaining authors declare no competing interests.

## Acknowledgements

This study was supported by JST, PRESTO Grant Number JPMJPR23I1, Japan, and JST Moonshot R&D Grant Number JPMJMS2012, Japan. The authors thank Shoko Tonomoto, Aya Kamiya, and Sayoko Ishii for their assistance.

## Notes

### Summary of Updates

In this revision, we improved the overall clarity and quality of the manuscript by thoroughly correcting grammatical issues and refining the scientific writing style. We also revised figure captions to ensure they are consistent with academic conventions and clearly describe the associated content. Additionally, we incorporated relevant references to support key claims, particularly those related to inter-subject variability in EEG-based decoding performance.

