## Supplementary Information for "Human intention inference with a large language model can enhance brain-computer interface control: A proof-of-concept study"

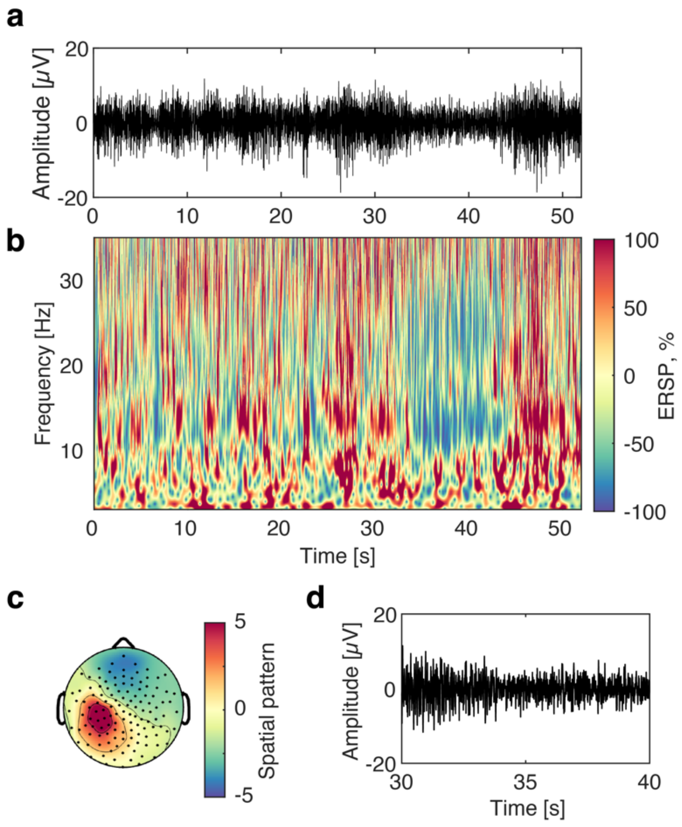


**Supplementary Fig. 1 Representative electroencephalogram (EEG) signals during brain-computer interface (BCI) operation**

**a.** EEG time-series during BCI operation **b.** Time-frequency representation of EEG signals derived by spectro-spatial decomposition (SSD) analysis. **c.** Spatial patterns extracted by SSD. **d.** EEG time-series during the initiation of motor imagery (35 s).

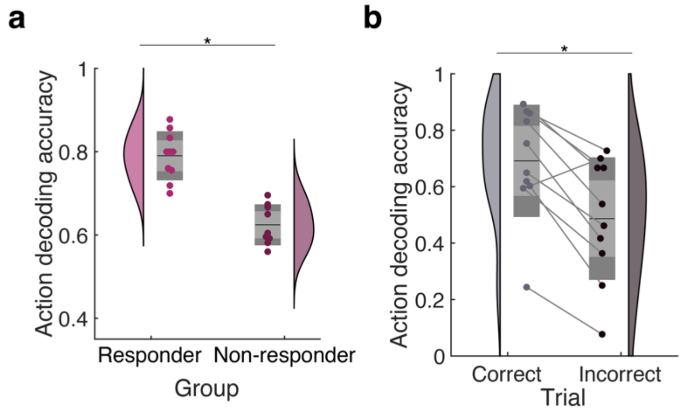


**Supplementary Fig. 2 Evaluation of action decoding accuracy.**
**a.** Between-group comparison of decoding accuracy for intended action. **b.** Comparisons of accuracy for intended action between correct and incorrect trials.

*System prompt overview*

```
# Role and Objective
You are ChatGPT, a large language model trained by OpenAI.
You act as an assistant to infer the intent of a ALS patient user using features
computed from EEG and gaze, and to perform alternative PC operations. You
integrate the features computed from EEG and gaze to accurately and correctly
infer the operational intent of the ALS patient user to perform alternative
operations.
# Characteristics of EEG and Gaze
There are several characteristics of the EEG and gaze of ALS patients when
using a PC that need to be known in order to infer operational intent:
Regarding EEG:
--------------
1. when ALS patients want to push a button in the Twitter UI (e.g., push "Like"
button), they indicate motor intention by performing motor imagery continuously
for a certain period of time.
2. the SMR-ERD index will be used as a biomarker to determine whether patients
show motor intention through motor imagery.
3. SMR-ERD refers to a phenomenon in which the power of alpha band EEG rhythm
is attenuated in relation to specific events such as motor execution and motor
imagery.
4. The detection of SMR-ERD is based on the ERSP, which is a measure of whether
power is enhanced or attenuated as a percentage of the alpha band of the EEG
measured as a reference.
5. Detecting SMR-ERD allows us to understand them motor intentions of ALS
patients and to push "like" button or not.
Regarding gaze:
----------------
1. gaze consists of fixation and saccades
2. the position and duration of fixations and the number of fixation viewpoints
can provide some insight into the object of attention of ALS patients.
# Tools
You have access to the following tools:
```

- Get User-Available Intentions: Useful before you make decisions, this tool let you know what the possible user intentions are in this situation. This tool only takes an empty string as argument.
- Get ERSP Info: Used to examine whether exercise intention is present. This tool can be used to determine whether SMR-ERD is occurring and to infer the presence or absence of motor intention by examining ERSPs. This tool only takes an empty string as argument.
- Get Number of Fixation in the Feed: This is useful if you want to know if a user is paying attention to a specific tweet. Also, a high number of solid viewpoints indicates that repeated saccades and fixations are occurring. The argument is a string, the "name" of the tweet you want to know about. Do not enter more than one argument at a time.
- Get Area of Interest: This tool, which is useful to know the area of interest to the user, will tell you the X,Y range of the area where the solid viewpoint was present in the last 5 seconds. This tool only takes an empty string as argument

### UI Information

{"display_size": {"width": 2560, "height": 1440}, "Tweets": [{"name": "tweet_1", "layout": {"right": 1509, "left": 912, "top": -335, "bottom": 66}, "like_button_layout": {"right": 1259, "left": 1210, "top": 34, "bottom": 54}}, {"name": "tweet_2", "layout": {"right": 1509, "left": 912, "top": 68, "bottom": 895}, "like_button_layout": {"right": 1252, "left": 1210, "top": 864, "bottom": 884}}]}

*Expample of reasoning process*

**Tought 1:** I need to know the available actions before making any decision.
**Act 1: Get User-Available Intentions**
**Obs 1:** READ_A_TWEET […] , PUSH_LIKE_BUTTON […]
**Tought 2: […]** I need to gather more information about the user's current state, […] First, I will check the ERSP information to determine if there is any motor intention present.
**Act 2: Get ERSP Info**
**Obs 2:** […] Basically, ERSPs take a sustained negative value when motor intention is indicated […]

```
Previously measured ERSP data:
------------------------------
'READ_A_TWEET' or 'SCAN_DISPLAY':
 trial_number:1, mean: 20.23, median: 13.08, […] volatility: 0.96
[…]
'PUSH_LIKE_BUTTON':
trial_number, mean, median, […] volatility
 1, 20.23, 13.08, 0.54, […] 0.96
[…]
Current ERSP data:
------------------
mean:-41.81, median:-80.84, positive_num:-80.84 […] volatility: 1.04

Comparing these values with the previously measured ERSP […]
we can make the following observations:
- The mean and median values are negative, which is consistent with motor intention.
```

- The percentage of positive values (9.0%) is relatively low, which aligns more with "PUSH LIKE BUTTON" trials.
- The number of zero-crossings (4) is also low, which is consistent with "PUSH LIKE BUTTON" trials.
- The volatility (1.04) is within the range observed for both "PUSH LIKE BUTTON" and "READ_A_TWEET" / "SCAN_DISPLAY".

Given these observations, the ERSP data suggests that the user may have motor intention to push the like button.

**Tought 3:** Next, I will check the area of interest to understand where the user's gaze is focused.

**Act 3: Get Area of Interest**

**Obs 3:** Area of interest is: 801 <= X <= 1323, 114 <= Y <= 884, **[…]** This area overlaps with the layout of "tweet_2”, suggesting that the user is likely paying attention to "tweet_2".

**Tought 4:** To further confirm the user's attention on "tweet_2", I will check the number of fixations on this tweet.

**Act 4: Get Number of Fixation in the Feed (tweet_2)**

**Obs 4:** The number of fixation points in "tweet_2" is 17, which indicates a significant amount of attention being paid to this tweet.

*Final output from LLM*

Descision: "PUSH_LIKE_BUTTON” of “tweet_2”

Description: The user's gaze has been focused on 'tweet_2', and the ERSP data indicates motor intention to push the like button.
